# The early local and systemic Type I interferon responses to ultraviolet B light exposure are cGAS dependent

**DOI:** 10.1101/835132

**Authors:** Sladjana Skopelja-Gardner, Jie An, Joyce Tai, Lena Tanaka, Xizhang Sun, Payton Hermanson, Rebecca Baum, Masaoki Kawasumi, Richard Green, Michael Gale, Andrea Kalus, Victoria P. Werth, Keith B. Elkon

**Author notes:** **Corresponding Author: Keith B. Elkon**, Division of Rheumatology, University of Washington, 750 Republican Street E531, Seattle, Washington 98109, USA.

## Abstract

Most systemic lupus erythematosus (SLE) patients are photosensitive and ultraviolet B light (UVB) exposure worsens cutaneous disease and precipitates systemic flares of disease. The pathogenic link between skin disease and systemic exacerbations in SLE remains elusive. In an acute model of UVB-triggered inflammation, we observed that a single UV exposure triggered a striking IFN-I signature not only in the skin, but also in the blood and kidneys. The early IFN-I signature was significantly higher in female compared to male mice. The early IFN-I response in the skin was almost entirely, and in the blood partly, dependent on the presence of cGAS, as was skin inflammatory cell infiltration. Inhibition of cGAMP hydrolysis augmented the UVB-triggered IFN-I response. UVB skin exposure leads to cGAS-activation and both local and systemic IFN-I signature and could contribute to acute flares of disease in susceptible subjects such as patients with SLE.

## Introduction

Sensitivity to ultraviolet B (UVB) light affects ~70% of systemic lupus erythematosus (SLE) patients and manifests as localized skin disease, cutaneous lupus erythematosus (CLE). However, acute UV exposure also instigates systemic disease flares^1,2^. Why SLE patients have such a high frequency of photosensitivity and how sunlight exposure leads to systemic disease exacerbations in SLE remain largely unknown. The chronically elevated expression of Type-I Interferon (IFN-I) and IFN-I simulated genes (ISG or IFN signature), in peripheral blood cells^3–5^, lesional and non-lesional skin^6,7^, as well as cells extracted from kidney biopsies^7,8^ provide clues to disease pathogenesis. *In vitro* studies of human keratinocytes have shown that exposure of cells to UVB light can induce an IFN-I response^9,10^. In this study, we ask whether acute exposure to UV light *in vivo* can trigger both a local and a systemic IFN-I signature. We also address differences in the response in female and male mice, due to the overwhelming bias of SLE for females (9:1 female to male ratio)^11^ and the recently reported sex-dependent differences in skin gene expression^12^.

In the only mechanistic *in vivo* study of UVB light-triggered ISG expression to date, we showed that repeated irradiation with a low dose of UVB light triggered local ISG expression via a stimulator of interferon (IFN) genes (STING)-dependent mechanism independent of pDC recruitment^13^. Canonically, STING has been thought to act primarily as a cytosolic DNA sensing adaptor protein downstream of cyclic GMP-AMP (cGAMP) synthase (cGAS), leading to production of IFNβ and inflammatory cytokines. However, recent studies have shown that other DNA (AIM2, IFI16) as well as RNA sensors (RIG-I-MAVS) signal through or cooperate with STING to drive IFN-I production in viral immune responses^14,15^. Involvement of the canonical cGAS-STING DNA sensing pathway in UV light-triggered IFN-I and inflammatory responses in the skin has not previously been examined.

Using a single dose of UVB light rather than repetitive stimulation (which is confounded by overlapping damage and repair responses), we first demonstrated that acute exposure to UV light triggers a strong IFN-I response in both normal mouse and human skin. Remarkably, UV light stimulated not only a local but also a systemic IFN-I response, in the blood and the kidneys. We found that the DNA sensor cGAS was required for the early local and systemic IFN-I signatures. Moreover, inhibiting the degradation of extracellular cGAMP by a chemical inhibitor of the cGAMP hydrolase ectonucleotide pyrophosphatase phosphodiesterase 1 (ENPP1) enhanced the magnitude of the IFN-I response.

## Results

### A single exposure to UVB light triggers an early cutaneous IFN-I response exaggerated in female mice

*In vitro* and *ex vivo* studies reported that UVB and UVC light-mediate cell damage induced IFN-I in keratinocytes and other cell types^10,16^. We and others observed that repeated exposure to low doses of UVB light (100mJ/cm^2^ per day for 5 days) caused a modest upregulation in ISG expression in the skin of normal mice^13,17^. However, this subacute model generates cycles of inflammation and resolution, which complicates understanding as to whether IFN-I reflects immediate injury or a wound repair response. Here, we first addressed whether a single dose (500mJ/cm^2^ in all experiments) of UVB light *in vivo* affects IFN-I production in C57BL/6J (B6) mice, a strain shown to best mirror cutaneous changes to UVB in human skin^18^. This, or higher doses have been widely used in the B6 mice and defined as 2 minimal inflammatory doses^9,19^.

Gene expression analyses of skin at different times (6, 24, and 48 hours) following exposure to a single dose of UVB light demonstrated a striking increase (~10-fold) in cutaneous ISG mRNA levels: *Irf7, Ifit1, Ifit3, Ifi44, Isg15, Isg20*, and *Mx1*, compared to non-UV exposed baseline (Fig. 1A). The mRNA expression levels of the majority of the tested ISG remained above baseline (1.8-16 fold induction) 48hr after exposure to UVB light, demonstrating that UVB-triggered IFN-I signaling persisted for several days (Fig. 1A). Similar to a previous report using *ex vivo* samples^16^, we detected a modest (~2-fold) and transient (6 hr) IFN-κ expression in the skin (not shown). Early induction in cutaneous expression of all the tested ISG was strikingly higher in female than in male mice, and the greater fold-induction of some ISG in female skin persisted at 24hr (*Ifit3 and Ifif44)* and 48hr (*Ifit3* and *Irf7)* after UVB exposure (Fig. 1A). Skin IFN scores (Fig. 1B), i.e. composite scores of the relative expression of each gene at specific time points relative to the baseline levels prior to UVB injury^20^, were higher in female compared to age matched male mice (mean IFN scores in female vs. male skin: 6hr = 8.56 vs. 1.0, 24hr = 8.7 vs. 3.6, 48hr = 4.9 vs. 2, Fig. 1B). These data demonstrate that females exhibit a markedly exaggerated early cutaneous IFN-I response to UVB injury (as much as 10-fold compared to males at 6hr). In light of these findings, all subsequent experiments were performed using female mice.

**Figure 1:**
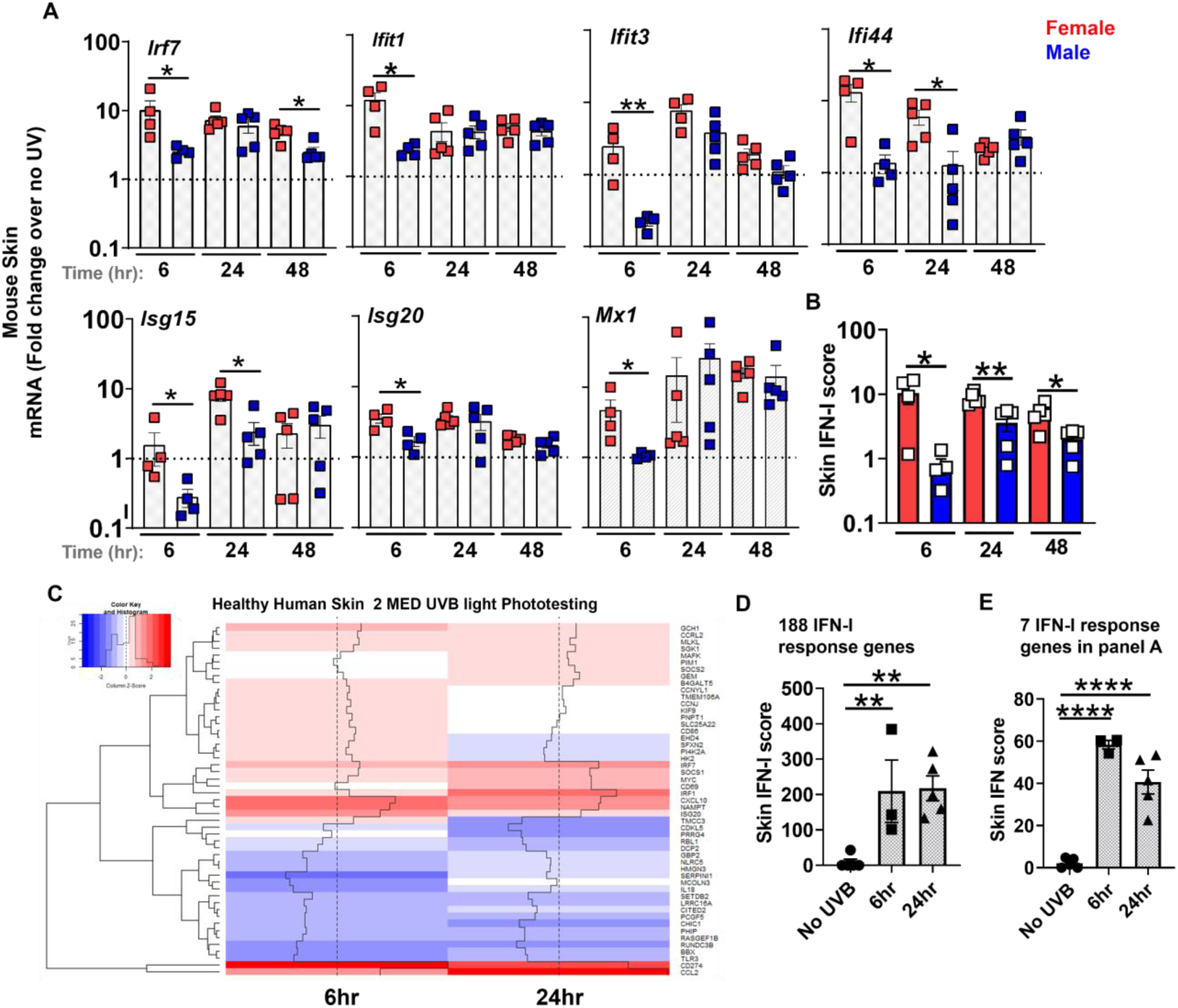
A single exposure of skin to UVB light triggers an early ~10-fold induction of type I IFN-stimulated genes (ISG) in female mice and induces interferon gene expression in human skin. Age-matched male and female B6 mice were shaved dorsally and the whole back was exposed to one dose of UVB light (500mJ/cm^2^). At 6, 24, and 48 hours after UV irradiation, (**A**) fold induction in the expression of type I IFN stimulated genes (ISG) in the skin: *Irf7*, *Ifit1*, *Ifit3*, *Ifi44*, *Isg15*, *Isg20*, and *Mx1* was determined relative to baseline, i.e. non-irradiated skin. (**B**) Skin IFN scores for female and male B6 mice at 6, 24, and 48 hr after UVB light irradiation were calculated as sum of normalized expression levels of the same 7 ISG as discussed in Methods. (**C-E**) IFN-I response was evaluated in healthy human volunteers 6 and 24hr after UV exposure (2 MED UVB) and the ISG most differentially expressed shown in the heatmap (C). IFN scores were derived based on (**D**) 188 published IFN-I response genes ^8^ or (**E**) 7 ISGs used in mouse studies (B). Statistical significance was determined by (A-B) Student’s t-test (*p<0.05, **p<0.01) or (D-E) one-way ANOVA (**p<0.01, ****p<0.0001).

### A single exposure to UVB light also induces an interferon signature in human skin

To test the relevance of our acute model of UV light-triggered IFN-I response in human skin, we exposed healthy volunteers (n=5, female) to a single dose of UV light (2 MED for UVB) and performed RNA sequencing on skin biopsies from unexposed skin and skin collected 6hr and 24hr after UV exposure. Using a published set of IFN-I response genes^8^, we demonstrated that a single UV exposure triggered an IFN-I signature in healthy skin as early as 6hr (Fig. 1C-D). Analogous to the murine studies, we observed a prominent IFN-I signature after UV exposure even when only the 7 genes investigated in the mouse skin were used to derive the IFN-I scores (Fig. 1E). Notably, the IFN-I scores derived from the 7 genes made up ~30% of the total IFN-score generated from the entire gene set (188 IFN-I response genes, Fig. 1D-E), suggesting that the 7 ISGs investigated in the murine studies are a reasonable representation of the overall IFN-I response in the human skin. These data for the first time demonstrate that UV light stimulates an IFN-I response in the human skin *in vivo.* Although it is well established that most SLE patients demonstrate an IFN-I signature in affected and unaffected skin^7,21^, the genesis of ISGs in the skin is unknown. Our findings suggest UV light exposure as a potentially important source of IFN-I activation in the skin in humans as well as mice.

### Skin exposure to UVB light triggers a systemic IFN-I response

The presence of an IFN signature in SLE patients was first detected in peripheral blood mononuclear cells (PBMC) in 2003^3–5^. More recently, the IFN signature in PBMC was reported in patients with cutaneous lupus^22^ and the IFN signature in SLE skin has been linked to a similar signature in lupus nephritis^7,8^. A key question in SLE is whether the IFN signature detected in PBMC is generated in blood or in tissues and, if in tissues, which tissues are responsible? To examine the systemic effects of UV light, we exposed mice to the single dose of UVB light and quantified ISG expression in circulating blood cells. As shown in Fig. 2A, UVB light-mediated skin injury triggered a striking upregulation in ISG mRNA levels in the peripheral blood cells (6-24hr). Compared to the skin, where ISG expression persisted 24hr after exposure to UVB (Fig. 1), the kinetics of ISG expression in the blood varied. Expression levels of most genes peaked early, while others were higher at 24hr (*Isg15* and *Isg20*) after UVB injury (Fig. 2A), possibly reflecting a second wave of injury, infiltration of immune cells, different DAMPs, or other mechanisms. The cumulative IFN scores reflect rapid induction in IFN signature in blood cells following skin exposure to UV light (Fig. 2B). The early IFN response in the blood was accompanied by increased IFNβ concentration in the plasma 6hr after exposure to UVB light (Fig. 2C). Of relevance to human SLE, pre-treatment of mice with hydroxychloroquine (HCQ), a TLR and cGAS antagonist^23^ that has demonstrated efficacy in treating cutaneous lupus in patients with an IFN signature^24^, significantly decreased the early IFN-I scores in both the skin and the blood after exposure to UVB light (Fig. 2D).

**Figure 2:**
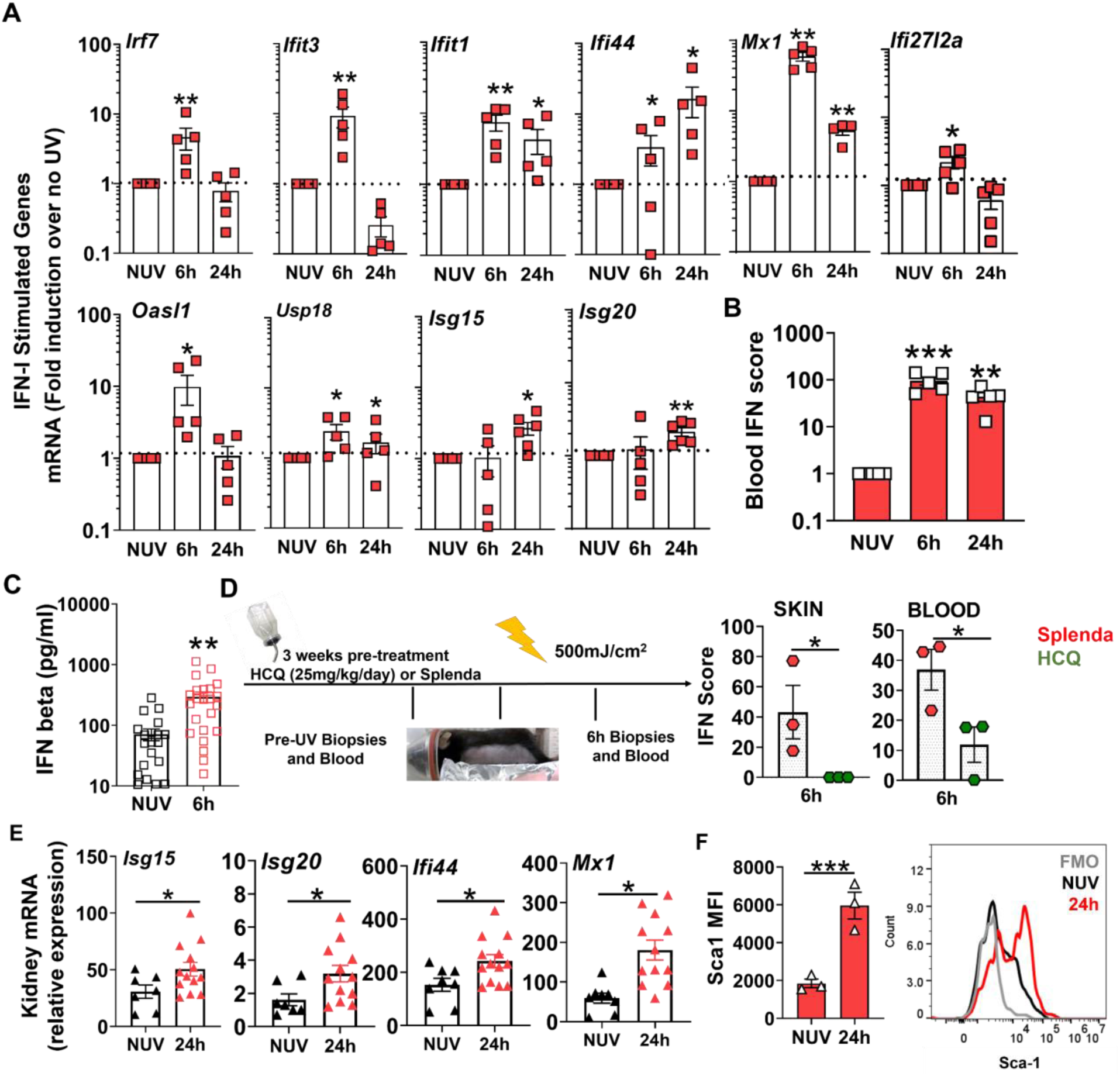
Acute skin exposure to UVB light triggers a systemic IFN-I response. Female B6 mice were exposed to a single dose of UVB light as in Fig. 1, except that only half of the back was exposed (**A**) Fold induction in ISG mRNA levels in the peripheral blood cells 6 and 24hr after skin exposure to UVB light was determined relative to mRNA levels in the blood prior to UV. (**B**) Blood IFN scores were calculated as the sum of normalized expression levels of the 7 most highly expressed ISGs after UV exposure (*Mx1, Ifit1, Ifit3, Ifi44, Usp18, Oasl1*, and *Ifi27l2a)*. (**C**) IFNβ concentration in plasma prior to UV (No UV, NUV) and 6h after UV light exposure in B6 mice. (**D**) B6 mice were treated with hydroxychloroquine (HCQ, 25mg/kg/day) or Splenda (controls) for 3 weeks prior to UV irradiation. Blood and skin IFN scores for both treatment groups were determined for samples prior to and 6hr after UV exposure. (**E**) Relative expression of representative ISG transcripts in the perfused kidney tissues of B6 mice at baseline (no UV, NUV) or 24h after skin exposure to UVB light. (**F**) Flow cytometry analysis of Sca-1 expression, presented as mean fluorescence intensity (MFI), on B cells in perfused kidney tissue prior to (NUV) and 24h after UV exposure. Statistical significance was determined by Student’s t-test (*p<0.05, **p<0.01, ***p<0.001).

Since recent studies reported that presence of an IFN signature in keratinocytes associates with lupus nephritis, as well as with an IFN-I signature in kidneys^7,8^, we asked whether UV light exposure of the skin could impact inflammatory markers, especially ISG, in the kidneys. Indeed, 24hr after skin exposure to UVB light, we observed an increase in ISG expression levels in kidneys (Fig. 2E), which could not be attributed to circulating blood cells as the kidneys had been perfused prior to harvest. The increase in Sca1 on intra-renal B cells (Fig. 2F) confirmed IFN-I exposure of cells^25^ in the kidney. Whether the IFN-I signature is explained by the direct influence of IFNβ, release of DAMPs from the skin, or even tissue infiltration by IFNβ producing cells that have migrated from the blood after skin exposure to UVB remain to be determined. In summary, analysis of blood and kidneys reveal that UV exposure of skin can act as a source of the systemic IFN-I in SLE, including the kidney.

### cGAS is essential for the early cutaneous IFN-I response to UVB light

UV light is genotoxic, cytotoxic, and generates modified nucleic acids such as oxidized DNA^10^. Recently, in a repetitive UV exposure model, we observed that IFN-I and inflammatory cytokines induced by UVB light were markedly reduced in STING deficient mice^13^. Canonically, STING has been thought to act primarily as a cytosolic DNA sensing adaptor protein downstream of cGAS. Upon engaging the cGAS synthesis product 2’3’ cGAMP, STING undergoes a conformational change to bind TANK binding kinase 1 (TBK1) and enables the phosphorylation of interferon regulatory factor 3 (IRF3) as well as activation of IKKβ and NFκB, leading to production of IFN-I and inflammatory cytokines, respectively^14^. However, other DNA (AIM2, IFI16, DAI) as well as RNA sensors (RIG-I-MAVS) can signal through, or cooperate with, STING so as to drive IFN-I production following certain viral infections^14,15^. Whether skin inflammation provoked by UV light triggers DNA to activate the canonical cGAS-STING pathway *in vivo* is not known. We therefore compared the ISG mRNA expression levels 6 and 24 hours following a single UVB light exposure of age-matched female B6 (wild type, WT) or *cGAS*−/− B6 mice, which have previously be shown to have intact TLR and other sensor pathways^26^. Whereas no difference in baseline (no UVB) skin ISG mRNA levels was found between WT and *cGAS−/−* mice (not shown), we observed that cGAS deficiency almost completely abrogated the early (6hr) cutaneous ISG expression after exposure to UVB (Fig. 3A). The lack of early ISG induction in *cGAS−/−* mice was comparable to that of IFN-α/β receptor deficient controls (*Ifnar−/−*) (Fig. 3A). While ISG expression in the *cGAS*−/− skin was detected 24hr after exposure to UVB, the fold-increase in ISG levels was substantially lower compared to WT controls (45.7% (*Ifit1)*-84.1% (*Ifit3)* reduction) (Fig. 3A). However, ISG mRNA levels in *cGAS−/−* skin were significantly higher than in *Ifnar−/−* controls at this later time point (Fig. 3A). Cumulative skin IFN scores confirmed that cGAS was required for the early cutaneous IFN signature after UVB injury, as ISG expression was not increased in cGAS−/− mice at 6hr. While later IFN-I scores in cGAS-deficient mice were lower than those in WT skin, they were significantly higher than in the *Ifnar−/−* controls (Fig. 3B), indicating that cGAS-independent pathways contributed to the IFN signature in the skin exposed to UV light over time. At this later time point, a modest increase in some ISGs (*Ifit1*, *ifi44*, Fig. 3A) in *Ifnar−/−* mice is consistent with previous reports of IFN-independent activation of these particular genes through IRF3-mediated gene expression^27,28^. Together, these data confirm that the requirement for cGAS in the cutaneous IFN-I response to UVB is temporally regulated. cGAS is essential for the early activation of IFN-I signaling and the cGAS-STING pathway is the dominant but not sole contributor to IFN-I activation in the skin later (24hr) following exposure to UVB.

**Figure 3:**
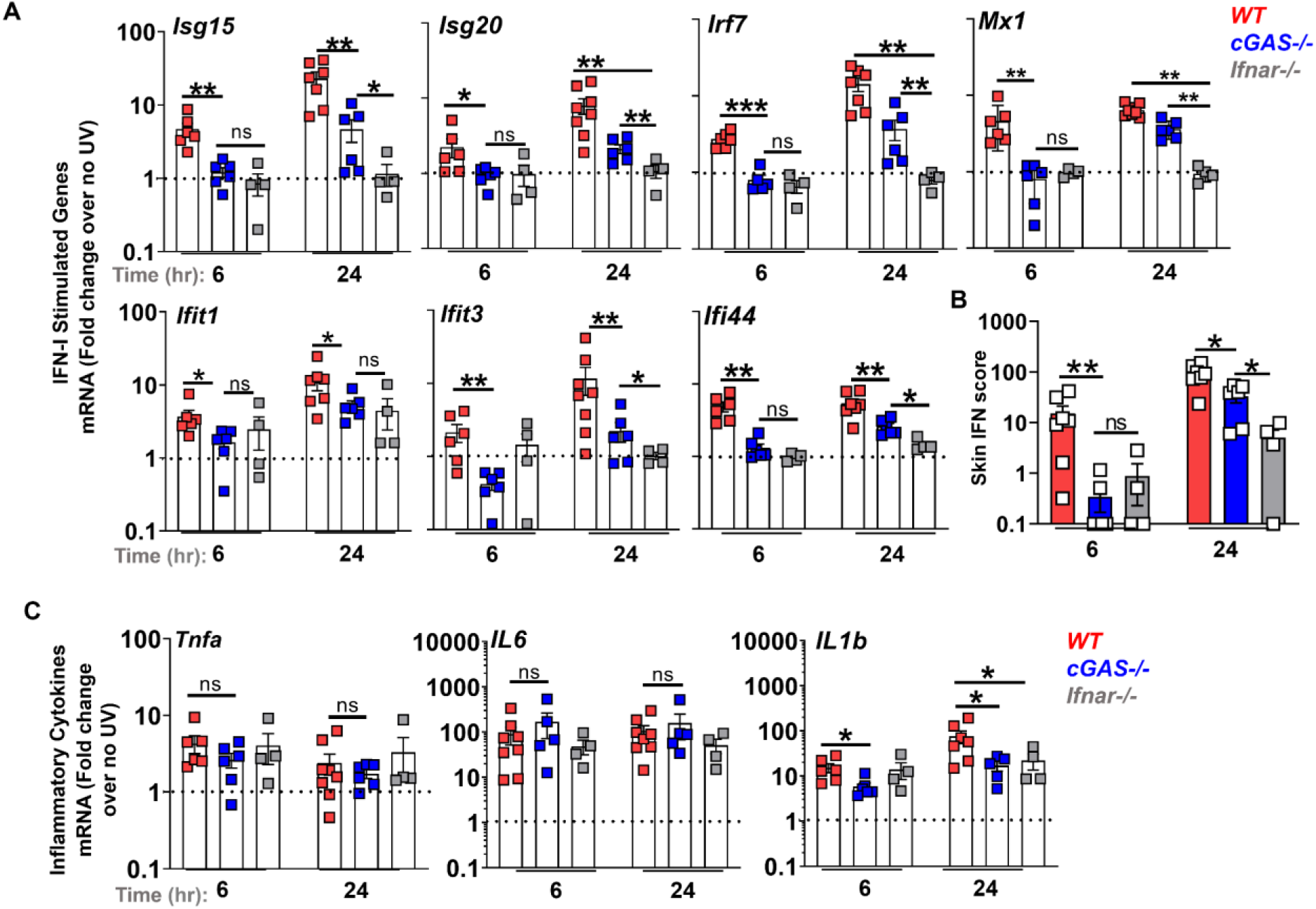
The early skin IFN response to UV light exposure is cGAS dependent. Age-matched female B6 (wild type, WT), *cGAS−/−*, and *Ifnar1−/−* mice were exposed to a single dose of UVB light as in Fig. 1, except that only half of the back was exposed. Skin biopsies were obtained prior to UVB light exposure and at 6 and 24h after irradiation. (**A**) Fold change in the expression of IFN-I stimulated genes (ISG) in the skin was determined relative to baseline, i.e. non-irradiated skin. (**B**) Skin IFN scores at 6 and 24 hours after UVB were calculated as sum of normalized expression levels of the same 7 ISG. (**C)** Fold-induction in the expression of inflammatory cytokines *Tnfa, Il-6*, and *Il1-b* were determined relative to baseline, i.e. non-irradiated skin. Statistical significance was determined by Student’s t-test (*p<0.05, **p<0.01, ***p<0.001, ns=not significant).

### TNF and IL-6 are stimulated via a cGAS-independent pathway following skin exposure to UV light

Since the cGAS-STING pathway has been implicated in the regulation of inflammatory cytokines through NFκB-mediated transcriptional activation of TNF and IL-6 or by activation of the inflammasome and generation of IL-1β^29^, we next asked whether UVB-triggered cGAS activation is necessary to induce these cytokines. We found that a single high dose of UVB light induced rapid expression (6hr) of TNF, IL-6, and IL-1β but that the extent of the TNF and IL-6 response was equivalent in WT and *cGAS−/−* mice (Fig. 3C). In contrast to TNF and IL-6, IL-1β mRNA levels were significantly lower in *cGAS−/−* skin both 6 and 24hr after UVB exposure (Fig. 3C). A significant reduction in IL-1β gene expression was also detected in IFNAR deficient mice, suggesting that intact IFN-I signaling is important for inflammasome activation following UV light injury (Fig. 3C). Together, our findings demonstrate that the DNA sensing cGAS–STING pathway is important for the generation of IFN-I and IL-1β, but not for stimulation of TNF and IL-6 following skin exposure to UV light.

### Extracellular cGAMP contributes to the cutaneous IFN-I and IL-1β response to UV light

Since extracellular export of the cGAS product, cGAMP, plays a role in the induction of IFN-I following radiation therapy^30^, we asked whether release of cGAMP following UVB radiation injury impacts IFN-I stimulation in the skin. To address this question, we inhibited the cell surface membrane enzyme ectonucleotide pyrophosphatase phosphodiesterase 1 (ENPP1), which hydrolyzes cGAMP^30,31^, shortly before skin exposure to UVB (Fig. 4A). The ENPP1 inhibitor, STF-1084^30^, led to a significantly greater induction in individual ISG expression as well as the cumulative IFN scores, compared to both the vehicle-treated skin exposed to UVB as well as to the unexposed skin treated with the ENPP1 inhibitor (Fig. 4B-C). Consistent with findings that cGAS was not required for UVB light-triggered TNF and IL6 production (Fig. 3C), inhibition of ENPP1 did not exaggerate cutaneous TNF and IL6 expression in response to UVB light injury (Fig. 4D). However, inhibition of cGAMP hydrolysis enhanced IL-1β expression in the skin after UV exposure (Fig. 4D), confirming that DNA sensing by cGAS contributes to inflammasome activation in UVB light-exposed skin (Fig. 3C). These data suggest that UVB exposure triggers cGAMP release which, in turn, amplifies the IFN signature and IL-1β, but not TNF and IL6, response.

**Figure 4:**
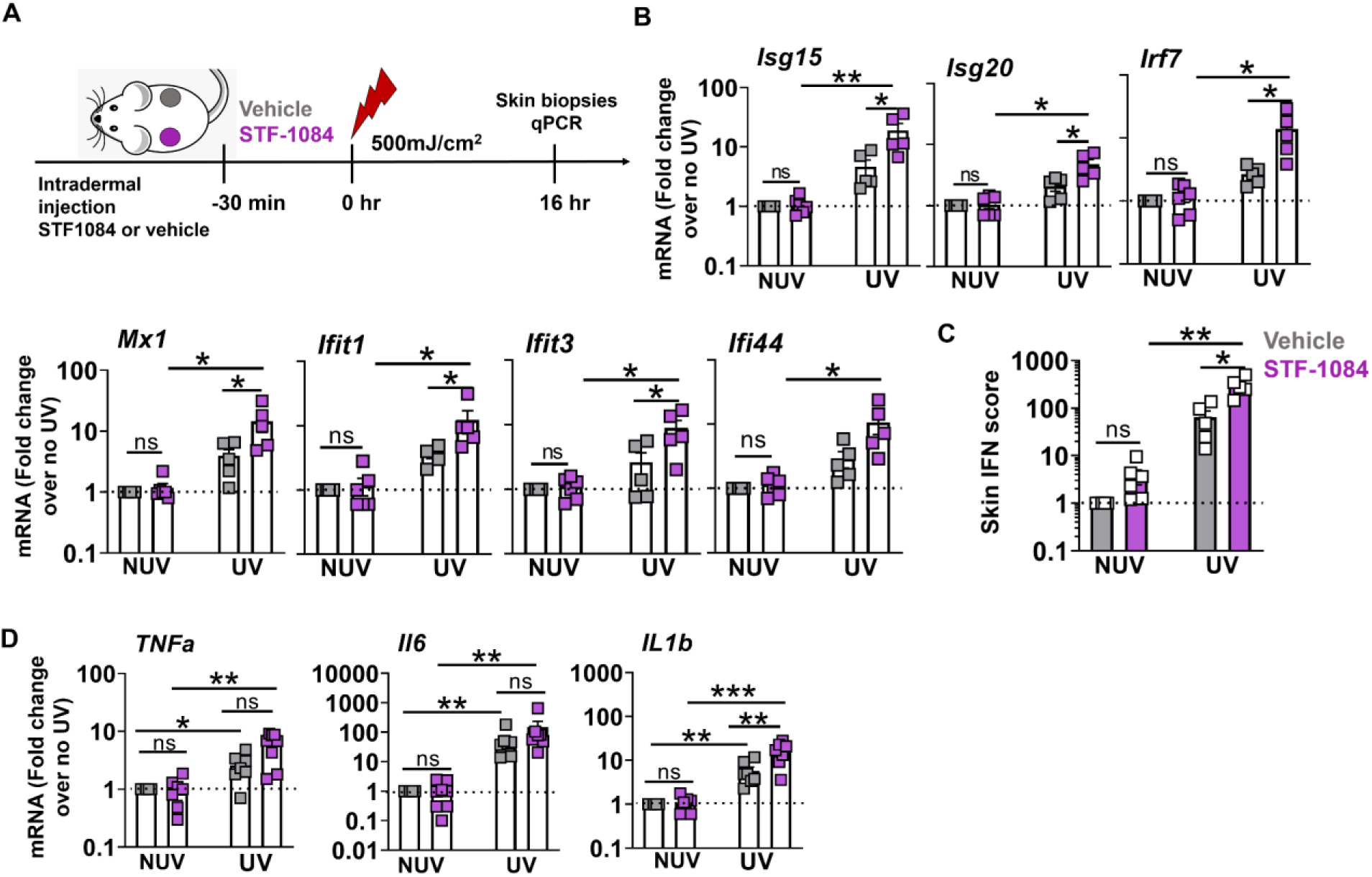
Extracellular cGAMP exaggerates the IFN-I, but not the IL-6 or TNF, response to UVB light. (**A**) B6 mice were shaved and injected intradermally with 100 µM ENPP1 inhibitor (STF1084) or Vehicle (PBS), 30 min prior to the exposure to UVB light as above. Skin was biopsied 16hr after UV exposure. (**B**) Fold-change in ISG expression was determined relative to vehicle-treated non-UVB exposed skin in three treatment groups: STF1084 without UVB, vehicle with UVB, and STF-1084 with UVB (n=5, 2 independent experiments). (**C**) Skin IFN scores in vehicle and STF-1084-treated UV exposed and non-exposed (NUV) skin were calculated as sum of normalized expression levels of the same 7 ISG as in Fig. 1. (**D**) Fold-induction in the expression of inflammatory cytokines *Tnfa, Il-6*, and *Il1-b* were determined relative to baseline, i.e. non-irradiated vehicle-treated skin. Statistical significance was determined by Student’s t-test (*p<0.05, **p<0.01, ns=not significant).

### cGAS mediates the systemic IFN-I response to UV exposure of the skin

Similar to the findings in the skin, early ISG expression in peripheral blood was largely abrogated in cGAS (by 45.1% (*Ifit1)* −95% (*Ifi44)* decrease) as well as IFNAR (~93% decrease) -deficient mice although the extent to which specific ISGs were affected differed (Fig. 5A). At 24hr after UVB injury, the expression levels of the majority of ISGs upregulated in WT mice were lower in the blood of *cGAS*−/− animals (by 57% (*Mx1)*-88% (*Usp18)*, Fig. 5A). To average the differences in the UV light-stimulated IFN signature in the 3 genotypes, we calculated the blood IFN scores using expression levels of 7 most-highly expressed ISGs. Overall, cGAS deficient mice demonstrated significantly lower blood IFN-I scores than their cGAS-sufficient counterparts, at both 6 and 24hr after UV light exposure (Fig. 5B). Peripheral blood cell IFN-I scores in *cGAS*−/− mice were higher than in the *Ifnar*−/− controls at 24hr post exposure indicating that other sensors besides cGAS also played a role at this time (Fig. 5B). Consistent with the lack of the IFN signature early in the blood of cGAS−/− mice (Fig. 3B), we could not detect an increase in the levels of circulating IFNβ (Fig. 5C) or upregulation of Sca-1 expression on cGAS deficient B cells (Fig. 5D). Therefore, in addition to the cutaneous response, a single exposure to UVB light stimulated an early cGAS-dependent IFN-I response in the blood.

**Figure 5:**
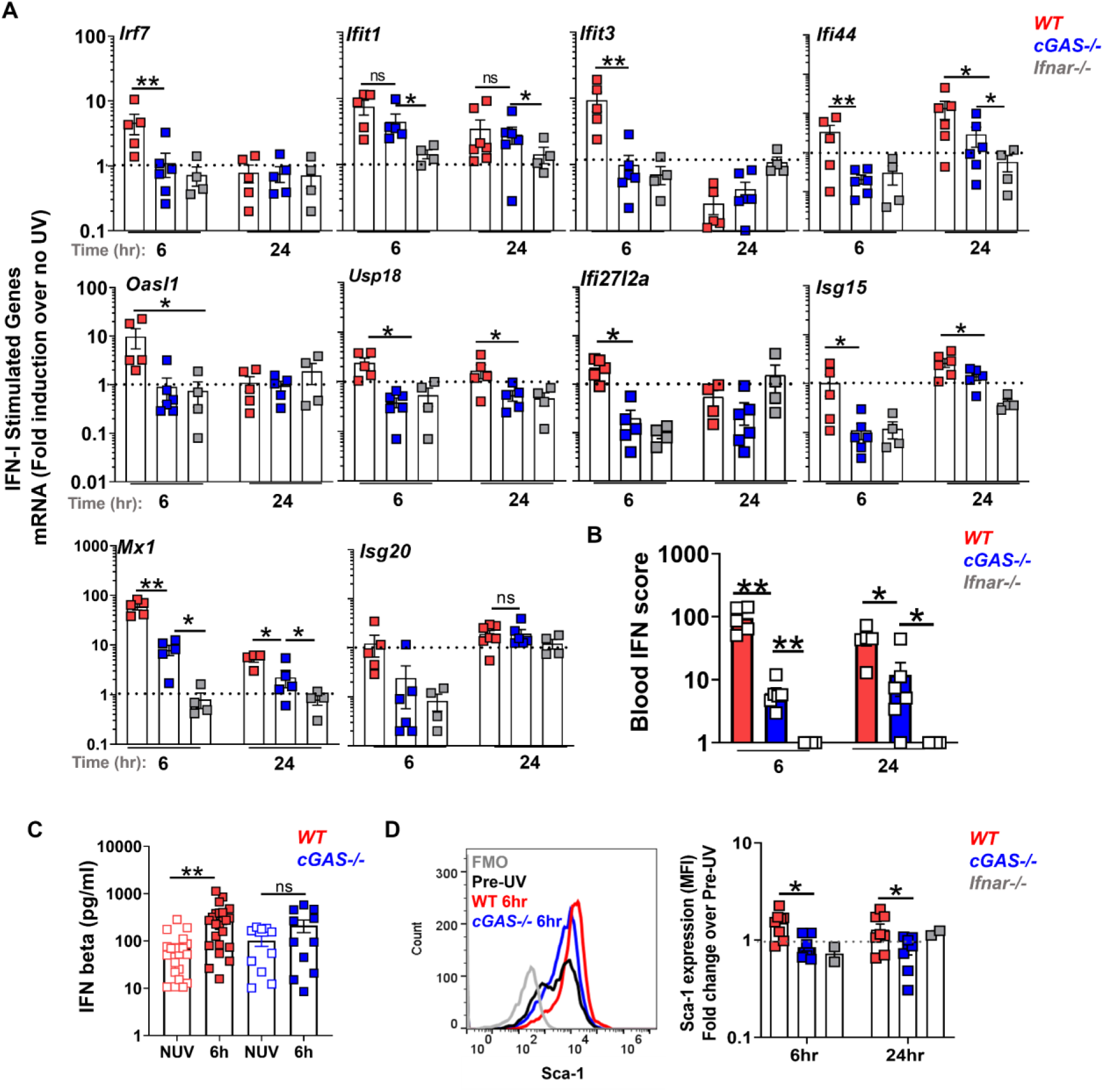
cGAS is required for the systemic IFN-I response to skin UV light exposure. Age-matched female B6 (wild type, WT), *cGAS−/−*, and *Ifnar1−/−* mice were exposed to a single dose of UVB light as in Fig. 2. (**A**) Fold induction in ISG mRNA levels in the peripheral blood cells 6 and 24hr after skin exposure to UVB light was determined relative to mRNA levels in the blood prior to UV. (**B**) Blood IFN scores for each genotype were calculated as the sum of normalized expression levels of the 7 most highly expressed ISGs after UV exposure (*Mx1, Ifit1, Ifit3, Ifi44, Usp18, Oasl1*, and *Ifi27l2a)*. (**C**) IFNβ concentration in plasma prior to UV (No UV, NUV) and 6h after UV light exposure in wild-type (WT) and *cGAS−/−* mice. (**D**) Flow cytometry analysis of Sca-1 expression on B cells in the blood of wild type (WT), *cGAS−/−*, and *Ifnar−/−* mice 6 and 24hr after UVB exposure, presented as fold change relative to non-irradiated skin cells. Representative histograms are shown for B cell population. Statistical significance was determined by Student’s t-test (*p<0.05, **p<0.01, ns=not significant).

### Absence of cGAS dampens the cellular inflammatory response in UV light-exposed skin

Analogous to inflammation triggered by infection^32^ and our observations of UVB light mediated sterile inflammation in the subacute model^13^, the acute response to UVB light in the skin was characterized by infiltration of neutrophils as well as inflammatory monocytes (CD11b+Ly6C^high^) at the site of injury (Fig. 6A-B). Besides innate immune cells, exposure to UVB triggered an increase in the number of CD8+ and γδ+ (Fig. 6C-D) but not CD4+ (not shown) T cells in the skin of WT mice. The number of both myeloid as well as T cell populations early (6hr) after UVB exposure was significantly reduced in the skin of cGAS-deficient mice (Fig. 6A-D). The diminished skin infiltration of inflammatory monocytes and CD8+ T cells in *cGAS*−/− mice was sustained 24hr after UVB exposure (Fig. 6B-C). Therefore, in addition to driving both the local and the systemic acute IFN-I response, cGAS-mediated DNA sensing also regulates the magnitude of the cellular infiltration into UV light-injured skin.

**Figure 6:**
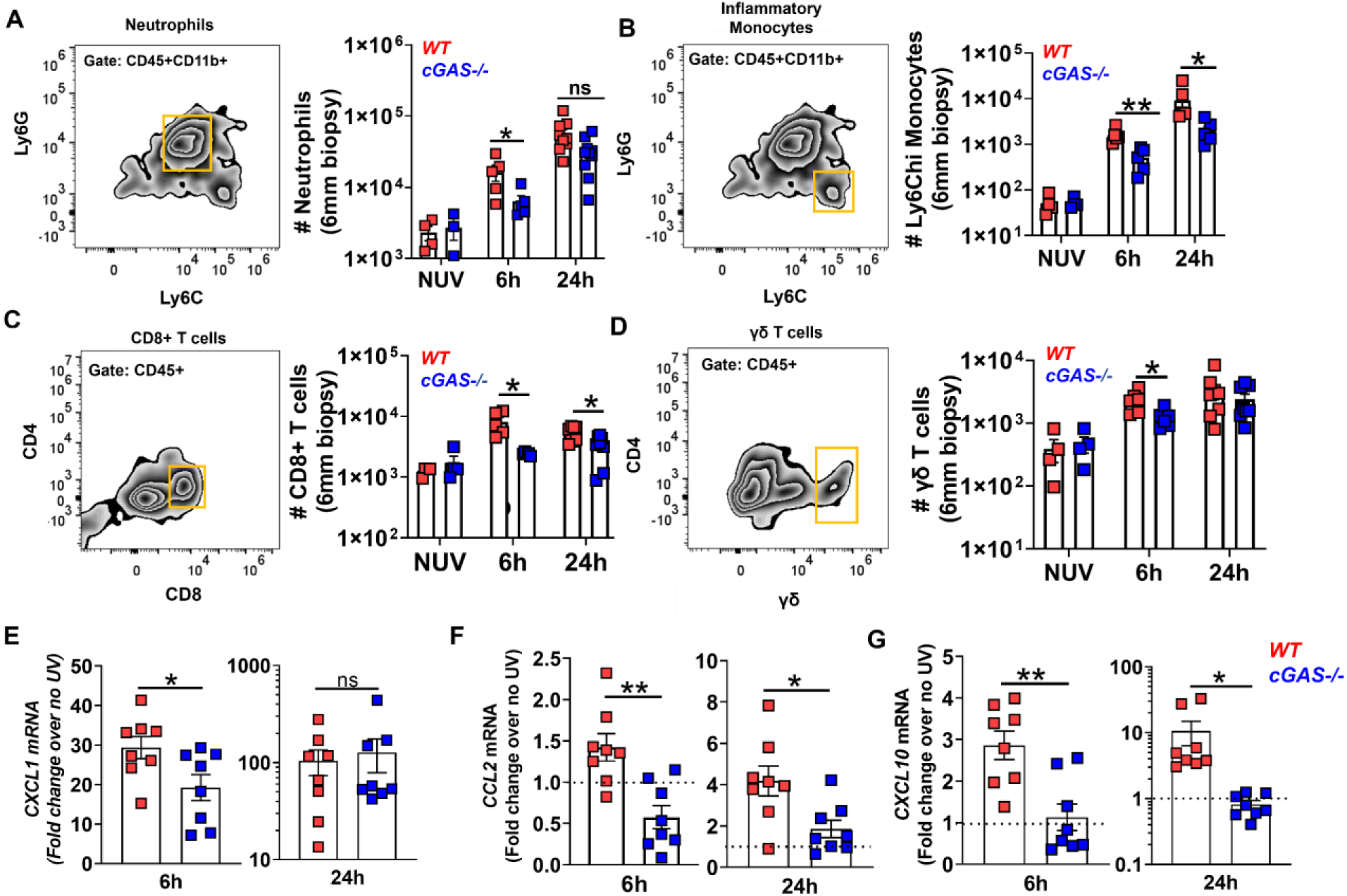
The inflammatory response to UVB light in the skin is diminished in the absence of cGAS. Age-matched female B6 (wild type, WT) and *cGAS−/−* mice were exposed to a single dose of UVB light as in Figs. 2–3. Flow cytometry analysis of skin was performed and the number of (**A**) neutrophils (CD45+CD11b+Ly6C^int^Ly6G^hi^), (**B**) inflammatory monocytes (CD45+CD11b+Ly6C^hi^Ly6G^neg^), (**C**) CD8+ T cells, and (**D**) γδ+ T cells determined based on total cell number per skin biopsy (6 mm). Skin biopsies from 2-4 mice were pooled for the 6h time point. (**E-G**) Gene expression levels of CXCL1 (E), CCL2 (F), and CXCL10 (G) in the skin were quantified by QPCR. Statistical significance was determined by Student’s t-test (*p<0.05, **p<0.01, ns=not significant).

To investigate whether diminished immune cell infiltration in the absence of cGAS was due to muted chemotactic responses following skin exposure to UV light, we evaluated the expression of the main neutrophil, monocyte, and T cell chemoattractants: CXCL1, CCL2, and CXCL10, respectively. Besides diminished production of IL-1β (Fig. 3C), we found reduced early expression of CXCL1 (Fig. 6E), both of which could contribute to lower neutrophil recruitment in the absence of cGAS. Lower cutaneous CCL2 expression in *cGAS−/−* mice (Fig. 6F) could explain the reduced monocyte numbers in skin. Finally, we found decreased CXCL10 expression in the skin of cGAS-deficient mice both early and 24hr after exposure to UVB (Fig. 6G), which likely contributes to the reduced CD8 T cell response to UV light^33–35^. Together, our data indicate that DNA sensing by cGAS contributes to innate and adaptive immune cell recruitment to UVB light exposed skin, possibly directly through IFN-I signaling and secondarily involving inflammasome activation and chemokine regulation.

## Discussion

In this *in vivo* study of the inflammatory effects of UVB light exposure, we report a number of novel observations. A single exposure to UVB light is sufficient to stimulate a robust IFN-I response in both murine and human skin. In mice, the early (6hr) IFN-I response is strikingly increased in females and the response is entirely cGAS dependent. Remarkably, not only does UVB exposure induce a local IFN-I signature, it also stimulates a systemic IFN-I response as determined by IFN signatures in the blood and kidney. The early blood IFN-I response was dependent on cGAS and was inhibited by hydroxychloroquine, which is consistent with our findings that aminoquinoline antimalarial drugs inhibit DNA activation of cGAS *in vitro* and *in vivo*^23,36^. Finally, we demonstrate that cGAS also impacts the cellular inflammatory response to UV light, as evidenced by reduced numbers of innate and adaptive immune cells in the UV light-exposed skin of cGAS deficient mice.

The detection of an IFN signature in the blood of SLE patients was a landmark discovery^3–5^, but where and how IFN-I is generated is not known. Here, we show that a single exposure of skin to UV light can generate an IFN signature in both the blood and kidneys of normal mice. Since ~70-85% of SLE patients are photosensitive and the skin has a vast surface area, it is plausible that IFN-I generated in the skin and released into the circulation is, at least in part, responsible for the blood IFN signature in SLE patients. That skin might be a source of systemic IFN-I in SLE is further supported by recent findings showing a 480-fold greater ISG expression in SLE non-lesional skin versus only a 7.8-fold in the SLE blood relative to healthy controls^37^. Intriguingly, individuals at-risk for developing SLE (ANA+, treatment naïve, ≤1 clinical criteria) also have a significant skin (28.7-fold increase) but lower blood IFN signature (2.2 fold increase) compared to healthy individuals^37^.

The significance of blood cell exposure to IFN-I following irradiation of skin with UVB is the potential for immune activation, especially in a genetically susceptible host. IFN-I can activate almost all cells in the immune system^38,39^ and Blanco et al^40^ showed that IFN-I in SLE blood causes differentiation of monocytes to DC with potent antigen presentation properties. Since many, but not all, clinical studies reveal an association between disease activity and the magnitude of the blood IFN-score^41,42^, the generation of an IFN signature in blood following UV exposure may well explain disease exacerbations associated with sunlight exposure^1,43^. Apart from diffusion of IFNβ, release of DAMPs, cell trafficking, or some combination of these factors may contribute to the blood IFN signature following exposure of skin to UVB.

In patients with cutaneous lupus, pDC are observed in lesional skin^21^. Furthermore, a Phase I/II clinical trial suggests that pDC depletion /inhibition (anti-BDCA2 therapy) ameliorates skin disease in cutaneous lupus patients^44^. Consistent with activation of the cGAS-STING pathway and failure to detect plasmacytoid dendritic cells (pDC) in the skin in our previous short-term UV study^13^, we observed that skin exposure to UVB light triggered an early increase in the circulating levels of IFNβ, in the absence of pDC (not shown). These apparently discordant findings could be explained by the fact that we are examining acute responses in healthy mouse skin versus immunopathological observations in lupus patients with chronic disease, circulating autoantibodies and a variety of other immunologic abnormalities. Whereas exposure of pDC to RNP containing immune complexes activates TLR7 resulting in high concentrations of IFNα in vitro^45^, it should be noted that pDC also produce IFNβ following cGAS-STING activation ^46^ and a recent analysis of the IFN signature in the skin of SLE patients identified a predominantly IFNβ driven gene response^47^. More detailed studies of the evolution of the IFN response following UVB exposure in normal and autoimmune prone strains will be needed to fully evaluate the role of different IFN-I species over time.

The recently discovered cytosolic DNA sensing cGAS-STING pathway has been implicated in generating IFN-I in response to viral infections ^48^, autoinflammatory and autoimmune disorders^49–51^, tumor cell-derived DNA, as well as DNA damage triggered by chemo- or radiation therapy in tumors^52,53^. Here, we observed that both local and systemic early production of IFN-I following skin exposure to UV light was almost entirely cGAS dependent. However, cGAS was not the sole contributor to the IFN-I response one day after UV exposure, demonstrating a temporal regulation of the mechanisms driving the IFN signature in response to UV light. Similar temporarily distinct roles for different DNA and RNA sensors have been described in virus infections. For example, following HSV1 infection, both cGAS and IFI16 were required for the early (6hr) IFNβ or TNF response but RIG-I was necessary for cytokine production at a later time point (16hr)^54^. Future studies will address whether other DNA and/or RNA sensing pathways that signal through or independently of STING contribute to the sustained IFN-I response after exposure to UV light.

UV irradiation triggers a number of biologic pathways in the skin including DNA damage, DNA repair, cell death, inflammation and resolution of skin damage^55,56^. Pathways of DNA damage and repair are complex and have been examined in response to UV light and ionizing radiation in the skin^57,58^. The inflammatory cytokine response in mouse skin includes increased expression of TNF, IL-6 and IL-1β^9,57^. Interestingly, we found that cGAS was required for stimulation of IFN-I, but not TNF or IL-6, in response to UV light. This selectivity suggests that the cGAS-STING pathway predominantly activates IRF-3 in UV light injured skin, while other pathways provide the main source of NF-κB signaling in this context. Two pathways stand out as possible contributors to this early inflammatory cytokine response as well as the later IFN-I response: TLR-3, which was responsible for stimulating TNF production in response to UV light-triggered RNA damage^9^, and ATM-IFI16-STING, which predominantly activated NF-kB and IL-6 production in response to etoposide-mediated DNA damage^14^. In addition to regulating the IFN-I response to UV light, we demonstrated that cGAS contributed to IL-1β expression in UV light exposed skin. While the relationship between IFN-I and IL-1β is complex, studies have shown that cGAMP can prime and activate the NLRP3 inflammasome^29,59^ likely explaining why IL-1β is lower in cGAS deficient compared to WT mice following exposure to UV light.

cGAMP is the cyclic dinucleotide messenger that engages STING to trigger transcription of IFN-I^48^. Recent studies of the cGAS-STING pathway in anti-tumor immunity have demonstrated that cGAMP can be released into the extracellular space where its lifespan is regulated by ENPP1, a cell surface phosphodiesterase that hydrolyzes cGAMP to AMP and GMP^30^. Interestingly, our findings that chemical inhibition of ENPP1 exaggerated the IFN-I response to UV light suggest that hydrolysis of extracellular cGAMP is an important regulatory mechanism following UV light stimulated inflammation. The disruption in cGAMP regulation also led to enhanced IL-1β, but not TNF or IL6, expression reinforcing the selectivity for the cGAS-STING-IRF3 pathway in shaping the inflammatory response to UV light. Whether a dysregulation in ENPP1 mediated hydrolysis of extracellular cGAMP could contribute to enhanced and persistent IFN signature in SLE skin, particularly in non-lesional areas, warrants investigation.

The ability of the cGAS-STING pathway to shape the inflammatory response via IFN-I signaling was recently identified in both spontaneous and radiation therapy-induced anti-tumor immunity^33,60^. Here, we found reduced infiltration of both the innate and adaptive immune cells in the UV light exposed skin in the absence of cGAS. The muted CD8+ T cell response we observed in the absence of cGAS was similar to the findings in the anti-tumor response, where cGAS-STING activation was required for the recruitment of tumor specific CD8+ T cells and their infiltration into the tumor environment^61–63^. These effects were largely attributed to the IFN-I response gene CXCL10, a recognized CD8+ T cell chemoattractant^33–35^. Consistent with these studies, we detected a reduction in CXCL10 expression in the skin of cGAS-deficient mice both early and 24hr after exposure to UV light. The role of cGAS in innate immune cell recruitment is less well established. Decreased neutrophil and inflammatory monocyte infiltration in UV light injured skin in cGAS deficient mice suggests an important role for the cGAS-STING pathway in shaping the innate immune inflammatory response. Diminished expression of neutrophil chemoattractants IL-1β and CXCL1 as well as monocyte chemokine CCL2 in the absence of cGAS provide a model by which DNA sensing influences innate chemotactic responses in UV light injured skin. While the effect of cGAS-STING signaling on monocyte recruitment in other inflammatory contexts is yet to be illuminated, cGAS-STING pathway appears to differentially influence neutrophil recruitment in infectious vs. sterile tissue injury^64,65^.

Besides providing novel mechanistic insights into the inflammatory response to UV light, our studies also introduced an important sex-dependent distinction in this response. The early skin IFN-I response to UV light was almost 10-fold higher in females compared to males which is of considerable interest considering the 9:1 female to male prevalence of SLE^11^. Whether this response can be explained by differential upregulation of the transcription factor, VGLL3, which is associated with heightened baseline expression of immune genes in the female skin^12^ remains to be determined. Interestingly, the greater ISG response in the skin in females is not restricted to provocation by UV light as females had higher ISG expression in response to HIV infection, possibly contributing to a greater protective role at the epithelial and mucosal surfaces^66^.

In summary, our findings indicate that exposure to UV light provokes a local and systemic IFN-I signature. IFN-I is stimulated initially via cGAS activation and this acute activation could mechanistically link UV light to flares of disease in SLE. In normal situations, acute inflammation is resolved through many reparative mechanisms that involve T-regulatory cells, Langerhans cells, and anti-inflammatory cytokines such as IL-10^17,56,67^. Whether SLE patients fail to control the inflammatory response to UVB in the skin and elsewhere is a critical question that will benefit from further understanding of the pathogenetic pathways involved.

## Methods

### Mice and UV irradiation

Male and female 12-16 week old C57BL/6 (B6, wild type), B6.*cGAS*−/−, or B6.*Ifnar−/−* mice were shaved dorsally. Mice were anesthetized with isoflurane and a single dose of UVB (500 mJ/cm^2^) was delivered either to the whole (figure 1) or half of the back (figures 2-5) using FS40T12/UVB bulbs (National Biological Corporation). The UVB light energy at the dorsal surface was measured with Photolight IL1400A radiometer with a SEL240/UVB detector (International Light Technologies). All animal experiments were approved by the Institutional Animal Care and Use Committee of the University of Washington, Seattle and guidelines for the care and use of animals during the research were followed. Mutant mice were kindly provided by Drs. Daniel Stetson and Michael Gale at the University of Washington.

### Detection of Interferon Stimulated Gene (ISG) Expression and IFNβ production

Skin biopsies (6mm) were performed prior to irradiation and at different time points after acute UVB injury: 6, 24, and 48 hours and tissue stored in RNA Later solution (Qiagen). Whole blood was collected prior to, 6hr, and 24hr following UVB light irradiation and red blood cells lysed. Kidneys were collected from mice following whole body cardiac perfusion with saline ^68^. RNA from skin and kidney was extracted by RNA Easy kit from Qiagen (Valencia, CA) and from blood cells using Quick RNA Miniprep (Zymogenetics). cDNA was synthesized using High Capacity cDNA synthesis kit (Applied Biosciences). ISG transcripts were selected according to previous studies of IFN response to UV and in *Trex*−/− animals ^13,36^ and quantified by real time quantitative PCR (qPCR) and normalized to *18S* (skin) or *Gapdh* (blood) transcript levels, using the following primers: *Isg15*, F:5′-AAGCAGCCAGAAGCAGACTC-3′ and R:5′- CACCAATCTTCTGGGCAATC-3′, *Isg20*, F:5’-TCACGGA CTACAGAACCCAAG-3’ and R: 5’ –TATCCTCCTTCAGGGCATTG-3’, IFN regulatory factor 7 (*Irf*7), F:5′- GTCTCGGCTTGTGCTTGTCT-3′ and R:5′-CCAGGTCCATGAGGAAGTGT-3′, *Mx1* F:5′- CCTCAGGCTAGATGGCAAG-3′ and R:5′ CCTCAGGCTAGATGGCAAG-3′*, Ifit1*, F:5′- TGCTGAGATG GACTGTGAGG-3′ and R:5′-CTCCACTTTCAGAGCCTTCG-3′*; Ifit3*, F:5’-TGGCCTACATAAAGCACCTAGATGG-3’ and R:5’-CGCAAACTTTTGGCAAACTTGTCT-3’; *Ifi44*, F:5’-AACTGACTGCTCGCAATAATGT-3’ and R:5’-GTAACACAGCAATGCCTCTTGT-3’, *Usp18*, F*: 5’-TTGGGCTCCTGAGGAAACC-3’ and* R*:5’-CGATGTTGTG TAAACCAACCAGA-3’, Oasl1*, F*: 5’-CAGGAGCTGTACGGCTTCC-3’ and* R*: 5’-CCTACCTTGAGTACCTTGAGCAC-3’. Ifi27l2a*, F*:5’-CTGTTTGGCTCTGCCATAGGAG-3’ and* R*: 5’-CCTAGGATGGCATTTGTTGATGTGG-3’, Gapdh*, F: 5’-TGGAAAGCTGTGGCGTGAT-3’ and R: 5’-TGCTTCACCACCTTCTTGAT-3’, and *18s*, F: 5’-AACTTTCGATGGTAGTCGCCGT-3’ and R: 5’-TCCTTGGATGTGGTAGCCGTTT-3’. Fold induction in ISG expression was determined using the standard formula 2 ^(-∆∆Ct)^ relative to baseline, i.e. skin prior to exposure to UVB. Mean IFN score was calculated as previously described ^22^: sum of normalized ISG expression levels; *i = each ISG;* SD = standard deviation.

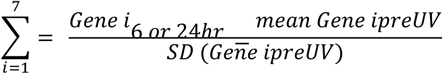

Interferon beta (IFNβ) protein levels in plasma were measured using Legendplex Mouse Inflammation Panel (Biolegend).

### Flow cytometry analysis of inflammation

Skin biopsies (1 6mm biopsy at 6hr and 5 6mm biopsies at 24hr) from female B6 and *cGAS−/−* mice were obtained prior to, 6hr, and 24hr after skin exposure to UVB light. Skin biopsies were processed as previously described ^13^. Briefly, the tissue was minced finely and digested with 0.28 units/ml Liberase TM (Roche) and 0.1 mg/ml of Deoxyribonuclease I (Worthington) in PBS with Ca^+2^ and Mg^+2^ for 60 minutes at 37°C with shaking. Cells were passed through a 0.7 μm strainer, resuspended in RPMI and counted. Cell were treated with Fc Block TruStain FcX (Biolegend) and stained in PBS + 1% BSA. Surface staining was performed using mouse–specific fluorescent antibodies purchased from Biolegend: i) Myeloid panel: CD45 APC.Cy7, CD11b PerCP.Cy5.5, Ly6C Violet 610, Ly6G FITC and ii) T cell panel: CD45 APC.Cy7, CD8 PerCP.Cy5.5, CD4 Pe.Cy7, γδ T PE. Samples were processed using CytoFLEX flow cytometer (Beckman Coulter) and data analyzed with FlowJo software v10 (Tree Star). Mean fluorescence intensities (MFI) of Sca-1 as well as percent positive populations were determined using fluorescence minus one (FMO) controls. Numbers of different cell populations were determined based on total counted cell numbers and normalized to the area of a single biopsy (6 mm).

### *In vivo* treatment with hydroxycloroquine (HCQ) and ENPP1 inhibitor

Mice (B6) were treated with HCQ (25 mg/kg/day) for 3 weeks administered orally in Splenda-sweetened water. Controls were treated with Splenda-sweetened water alone. Following 3 weeks of treatment, mice were exposed to UVB light as described above. Blood and skin taken prior to and 6hr after UV exposure were analyzed for ISG expression by qPCR and IFN scores derived, as described above. For treatment with ENPP1 inhibitor, mice were injected intradermally with 50µl 100 µM STF1084 (two injections per mouse) (generously provided by Yinling Li at Stanford University), as used by Carozza et al for intra-tumoral injections ^30^. After 30 minutes, mice were irradiated with a single dose of UVB light as described above. Skin biopsies were collected after 16hr and processed for RNA isolation and QPCR as above. Skin was analyzed for ISG expression under four different conditions: vehicle alone, STF-1084 alone, vehicle + UVB, and STF1084 + UVB. Fold change in ISG expression was determined relative to vehicle alone control skin. IFN scores were derived as described above.

### Human phototesting and skin RNA Sequencing

Healthy volunteers were recruited from the University of Washington (UW, Seattle WA) and Philadelphia Veterans Affairs Medical Center (Pennsylvania, PA). All individuals signed an informed consent in respective IRB-approved protocols (University of Washington; HSD number 50655). All methods were carried out in accordance with relevant guidelines and regulations for human participants at the University of Washington. The sun-protected skin of 5 female healthy controls was exposed to two minimal erythematous doses (MED) of UVB light on the dorsal surface of the lower arm. Solar Simulator radiation was generated using a 150 W xenon arc, single port Berger Solar Simulator (Solar Light, Model # 16S-150/300, Glenside, PA). UG-5 and WG 320/1inch thick glass filters was fitted to give a wavelength spectrum that includes wavelengths above 290nm-400nm. A 6-mm punch skin biopsy was collected from unexposed skin, and from skin 6hr and 24hr after UV exposure. Tissue was stored in RNA later and samples from PA shipped to UW. Following RNA isolation at UW, the RNA yield was 2-3 ug RNA (all samples had RIN >9.5). RNA sequencing was performed at Northwest Genomics Center at the University of Washington. cDNA libraries were prepared from 1 μg of total RNA using the TruSeq Stranded mRNA kit (Illumina, San Diego, CA) and the Sciclone NGSx Workstation (Perkin Elmer, Waltham, MA). Prior to cDNA library construction, ribosomal RNA was removed by means of poly-A enrichment. Each library was uniquely barcoded and subsequently amplified using a total of 13 cycles of PCR. Library concentrations were quantified using Qubit fluorometric quantitation (Life Technologies, Carlsbad, CA). Average fragment size and overall quality were evaluated with the DNA1000 assay on an Agilent 2100 Bioanalyzer. Each library was sequenced with paired-end 75 bp reads to a depth of 30 million reads on an Illumina HiSeq 1500 sequencer.

### RNAseq Data Processing and Analysis

Raw RNAseq data (Fastq files) were demultiplexed and checked for quality (FastQC version 0.11.3). Next rRNA was digitally removed using Bowtie2 (version 2.2.5). Sufficient host reads (~thirty million) were then mapped to the Human genome (GRCh37) using STAR (2.4.2a) and then converted into gene counts with ht-seq (0.6.0). Both the genome sequence (fasta) and gene transfer files (gtf) were obtained using Illumina’s igenomes website (https://support.illumina.com/sequencing/sequencing_software/igenome.html). Gene counts were then loaded in the R statistical programming language (version 3.2.1) and filtered by a row sum of fifty or more across all samples. Raw fastq files and count matrix were submitted to GEO and available under accession GSEXXX. Exploratory analysis and statistics were also run using R and bioconductor. The gene count matrix was normalized using EDGER and transformed to log counts per million (logCPM) using the voom through the limma bioconductor package (3.24.15). Statistical analysis (including differential expression) was performed using the limma package ^69^.

#### Interferon Z-score heatmap

Differential expression data was filtered for interferon genes and then plotted using the heatmap.2 function with the gplots bioconductor package in R ^70^.

#### IFN score analysis

IFN-I scores at 6 (n=3) and 24hr (n=5) after UV were derived using the established IFN-I response gene dataset (188 of 212 genes were detected ^8^. Relative expression of genes at baseline (prior to UV, n=5) and at different times after UV was normalized as described above and IFN scores generated using the same formula as for mouse studies, using either all 188 genes or just the 7 genes used in mouse studies.

### Statistical Analysis

Data were analyzed using GraphPad Prism 7 software (GraphPad Software Inc.) and presented as mean ± SEM. Statistical difference between two data groups was determined using Student’s *t*-test. One way-ANOVA was used to determine statistical significance between three groups in human UV exposure studies. *P* < 0.05 was considered significant.

## Acknowledgments

This work was supported by National Institutes of Health R21 AR072377-01 and ACR Rheumatology Research Foundation Grant to Keith Elkon; Veterans Affairs Merit Review [I01BX000706] to VPW and NIH/NIAMS Werth R01AR071653. The RNA Sequencing work was supported by an NIH grant for the University of Washington Interdisciplinary Center for Exposures, Diseases, Genomics & Environment, P30ES07033. We thank Tomas Mustelin, Jeffrey Ledbetter, Edward Clark, and Erika Noss for insightful discussion. We thank Lingyin Li at Stanford University for generously providing the ENPP1 inhibitor STF-1084.

The authors declare no competing financial interests.

## Author contributions

S.SG. and K.B.E. conceived the study and wrote the manuscript. S.SG., L.T., X.S., J.T., P.H., and R.B. conducted the experiments. S.SG. analyzed the data. J.A. and M.K. contributed to the conceptualization of the study and experimental design. M.G. and R.G. performed the analysis of RNA sequencing data. V.P.W., K.B.E., and A.K. conceptualized and executed human photo testing study. All authors critically reviewed and edited the manuscript.

